# Characterization of environmental effects on flowering and plant architecture in an everbearing strawberry F1-hybrid by meristem dissection and gene expression analysis

**DOI:** 10.1101/2022.06.12.495836

**Authors:** Samia Samad, Rodmar Rivero, Pruthvi Balachandra Kalyandurg, Ramesh Vetukuri, Ola M. Heide, Anita Sønsteby, Sammar Khalil

**Affiliations:** Department of Biosystems and technology, Swedish University of Agricultural Sciences, Alnarp, Sweden; NIBIO, Norwegian Institute of Bioeconomy Research, 1431 Ås, Norway; Department of Plant Breeding, Swedish University of Agricultural Sciences, Alnarp, Sweden; Faculty of Environmental Sciences and Natural Resource Management, Norwegian University of Life Sciences, Ås, Norway

**Keywords:** Everbearing strawberry, Floral induction, Floral initiation, Flowering, *Fragaria* x *ananassa* Duch., Photoperiod, Temperature

## Abstract

Floral transition in the cultivated everbearing strawberry is a hot topic because these genotypes flower perpetually and are difficult to maintain in a non-flowering state. However, it has rarely been studied using morphogenetic and molecular analyses simultaneously. We therefore examined morphogenetic effects and the activation of genes involved in floral induction and initiation in seedlings of an everbearing F1-hybrid. Seedlings were grown at 12, 19, and 26°C under 10-h SD and 20-h LD conditions. We observed a strong environmental influence on meristem development and a *FLOWERING LOCUS T1 (FaFT1)–SUPPRESSOR OF OVEREXPRESSION OF CONSTANS1 (FaSOC1)* pathway similar to that in the everbearing woodland strawberry. The everbearing cultivar showed typical features of a quantitative LD plant, flowering earlier under LD than SD conditions at all temperatures. We also found that floral induction is facilitated by *FaFT1* upregulation under LD conditions, while *FaSOC1* upregulation in the apex leads to photoperiod-independent floral initiation. Moreover, we confirmed the strawberry meristem identity gene *FRUITFULL* (*FaFUL)* can also be used as an early indicator of floral initiation in EB cultivars. This study also highlights the advantages of using seed-propagated F1-hybrids for genetic studies because are genetically identical, and not biased by a previous flowering history.

## 1. Introduction

The cultivated strawberry, *Fragaria × ananassa*, is a globally important fruit crop, so there is great interest in increasing its production in terms of quality, the quantity produced, and the length of its marketing season. Production can be increased by cultivating strawberries in protected environments such as greenhouses and tunnels and by using everbearing (EB) cultivars that flower perpetually [1–4]. Understanding the genetics of flowering will enable breeders to produce new cultivars and identify conditions that promote flowering and increase yields.

The transition from the vegetative to the reproductive stage is an important step in plant development that is regulated by both internal and external factors. It can be divided into four stages: induction, initiation, differentiation, and development. Induction occurs when the leaf perceives an environmental cue to flower, while initiation encompasses all of the physiological and morphological transformations that occur in the meristem after induction. The subsequent steps are differentiation, i.e., the formation of the floral organs, and anthesis, which involves the production of open flowers [5].

Flowering physiology in strawberry is governed by a strong and complex photoperiod x temperature interaction. The common June bearing or seasonal flowering (SF) cultivars are facultative short day (SD) plants [6,7] in which flowering is photoperiod-independent at low temperatures (≤12°C), requires SD conditions at intermediate temperatures (18°C – 21°C), and is repressed at high temperatures (>24°C) [7,8]. The exact critical temperatures and photoperiods for these responses may vary with genotype, growing conditions, and duration of SD treatment [3,9]. EB cultivars are also photoperiod-insensitive at low temperatures (<15°C) but become quantitative long day (LD) plants at intermediate temperatures (18-21°C) and obligatory LD plants above 24°C [2,10].

Strawberry plug plants obtained by clonal propagation are produced and maintained to preserve their genetic traits. This is a costly and laborious practice [11] that can cause uneven flowering due to variation in the environmental conditions perceived by the mother plants [4,10,12]. Furthermore, in commercial production, strawberry plants are raised under inducing conditions and then transferred to a greenhouse for flowering and fruiting when floral initiation is complete. However, several studies have shown that environmental factors can influence flower bud differentiation and subsequent development even in plants that have been induced and initiated flower buds [4,10–16]. This must be taken into account because initiating adequate numbers of inflorescences is of similar economic importance to producing runners for clonal propagation [11]. Detailed knowledge of environmental effects on meristem development is therefore needed to optimize these processes. For this reason, meristems are traditionally subjected to laborious microscopic evaluation of their floral development by so-called “flower-mapping” before the plants are transferred to greenhouses for flowering [11,15,17,18].

Although the physiology of flowering in the cultivated strawberry has been widely studied [2–5, 8–15], there have been comparatively few genetic studies on this topic [17,24–29]. The *Arabidopsis* photoperiodic pathway is conserved in the octoploid strawberry, but because the genetics of the cultivated strawberry are complex, most functional studies on this topic have been conducted in the closely related diploid wild strawberry *Fragaria vesca* [30–34]. *F. vesca* also has two different flowering types, namely seasonal flowering SD types and EB LD types, the latter of which only flower freely at high temperatures (27°C) under LD conditions [2]. The SF genotypes initiate flowers at low temperature (<12°C) independently of the photoperiod, are obligate SD plants at intermediate temperatures (12-16°C), and cannot flower at higher temperatures [2,35]. Conversely, flowering in the EB genotypes is promoted by LD conditions and temperatures above 18°C, and is delayed at low temperatures under SD conditions [30].

In the SF diploid strawberry, mRNA of the *CONSTANS* homologue *FvCO* accumulates in LD, leading to increased production of the stabilized CO protein and subsequent activation of *FLOWERING LOCUS T (FvFT1)*. Therefore, *FvFT1* is only upregulated in leaves under LD conditions [31,32,35,36]. The FT protein then moves to the shoot apical meristem (SAM), where all of the post-induction processes occur. In the apex, the FT protein activates *SUPPRESSOR OF OVEREXPRESSION OF CONSTANS1 (FvSOC1)* [30], which in turn activates *TERMINAL FLOWER1 (FvTFL1)* [30], a strong floral repressor [31]. Therefore, flowering is inhibited in LD in the diploid SF strawberry. Additionally, Rantanen et al. (2015) observed that *FvTFL1* is upregulated by high temperatures, explaining why SF *F. vesca* does not flower at high temperature. In EB genotypes, the same pathway exists but a 2-bp deletion in *FvTFL1* results in a non-functional repressor so that EB genotypes flower even at high temperature in LD but only marginally in SD. In both genotypes, the floral meristem identity genes *APETALA1* (*FvAP1)* and *FRUITFULL* (*FvFUL1)* are upregulated during floral initiation [30,31]. It was also shown that *FvSOC1* regulates photoperiod-dependent runner production in the axillary meristems via the action of *GIBBERELLIN20-oxidase4* (*FvGA20ox4*). However, runner production is promoted independently of *FvSOC1* at higher temperatures [37].

In contrast, a single locus governs the EB trait in the octoploid strawberry [38] but Castro et al. (2015) and Honjo et al. (2016) found that complex environmental (GxE) interactions made it difficult to isolate the EB gene [39,40]. Despite this, some flowering genes along the photoperiodic pathway have been characterized in the octoploid [17,24,25,29]. Nakajima et al. (2014) were the first to isolate *FT* and *TFL* genes in an SF cultivated strawberry (specifically, cv. ‘Tochiotome’). Nakano et al. (2015) and Koskela et al. (2016) studied *FaFT1* expression in the leaves of SF cultivars and showed that its daily oscillation matches that of *FvFT1*. Koskela et al. (2016) also functionally characterized *FaFT1* expression in the cultivars ‘Glima’ and ‘Elsanta’, revealing that it is expressed more strongly in LD than in SD, in accordance with the findings of Nakajima et al. (2014) and Nakano et al. (2015) for the Japanese cvs. ‘Tochiotome’ and ‘Nyoho’, respectively [24,25,29]. Moreover, two *FT*-like genes from the diploid strawberry, *FvFT2* and *FvFT3*, were isolated and overexpressed in the octoploid SF cultivar ‘Sveva’, where *FvFT2* promoted flowering while *FvFT3* promoted runner formation [26]. More recently, Liang, et al. (2022) analyzed the transcriptome of the SF cv. ‘Benihoppe’ and reported similar genes were activated during floral induction as that in the diploid strawberry [41]. The genetics of flowering in SF cultivars have thus been studied in some detail [17,24,27–29]. However, less is known about the genetics of flowering in EB cultivars.

Liang, et al. (2022) also showed that Knowledge of flowering genes can be used to rapidly identify the critical point of initiation [17,27,41]. We have therefore studied the relationship between environmental effects on the morphogenetic changes of the meristem and the expression of known flowering-related genes in the cultivated strawberry [24,25].

Here we present studies on the induction and initiation stages of flower development in the EB ‘Delizimmo’ cultivar conducted to clarify the flowering process in EB cultivars. Specifically, we investigated meristem development in both stages and correlated it to changes in the expression of known flowering genes. To ensure that the starting material in these studies had no prehistory of being induced [4,12], we used seed propagated F1-hybrids. This enabled us to compare the reliability of flower mapping and molecular analysis as practical tools for strawberry growers and breeders, and to determine how genes shape the flowering phenotype. Our results provide new insights into the relationship between the physiology and genetics of flowering that will be valuable for improving commercial strawberry production in many countries around the world.

## 2. Materials and Methods

### 2.1. Plant material, growing conditions and sampling

Seeds of the strawberry F1-hybrid cv. Delizzimo (ABZ Seeds, Bovenkarpel, The Netherlands) were sown on March 16 in sowing trays in a peat-based potting compost (Gartnerjord, LOG, Oslo, Norway) in a greenhouse at a minimum temperature of 24°C with a 10-h photoperiod at an irradiance of 150 μmol m^−2^ s^−1^. After 10 days (on March 26), when the seeds had germinated, the trays were moved into the daylight phytotron at the Norwegian University of Life Sciences at Ås, where they were exposed to a constant temperature of 26°C and a photoperiod of 10 h. On April 14, when the plants were well-rooted and had reached heights of 1 to 2 cm, they were potted in 9 cm plastic pots filled with granulated vermiculite, set on trolleys, and further grown at 26°C under 10-h SD conditions until April 30. The plants were then exposed for 5 weeks to constant temperatures of 12°C, 19°C, or 26°C with photoperiods of 10 or 20 h.

In the phytotron, the plants received natural daylight for 10 h per day (08.00-18.00 h). Whenever the photosynthetic photon flux (PPF) in the daylight compartments fell below approximately 150 μmol m^−2^ s^−1^ (as on cloudy days), an additional 125 μmol m^−2^ s^−1^ were automatically supplied using high-pressure metal halide lamps (400 W Philips HPI-T). Daylight extension to 20 h long days (LD) was achieved by applying low intensity illumination using 70 W incandescent lamps (c. 7 μmol m^−2^ s^−1^ PPF) such that the 4 h dark period was centered around midnight (22.00 h to 02.00 h). Plants subjected to short day (SD) treatment were kept in darkness from 18.00 h to 08.00 h. The plants thus received nearly the same daily light integral in both photoperiods. The plant trolleys were randomly positioned in the daylight rooms as a result of the every-day movements to and from the adjacent photoperiodic treatments rooms. Temperatures were controlled to within ±1.0°C and a water vapor pressure deficit of 530 Pa was maintained at all temperatures. Throughout the experimental period, the plants were irrigated daily to drip-off with a complete fertilizer solution consisting of a 1:1 (w:w) mixture of Kristalon™ (9-11-30% NPK + micronutrients) and Yaraliva™ (N 15.5% and Ca 19%) both from Yara International (Oslo, Norway) with an electrical conductivity (EC) of 1.5 mS cm^-1^. Three plants per treatment were harvested for phenotyping during the pre-treatment (on April 16), and in weeks 0, 3, and 5 of the experiment (Table 1). The pre-treatment was necessary to show that the plant was in a completely vegetative state prior to the start of the treatment and to establish a baseline for the genetic analysis. There have been reports [10,12,42] that EB plants can be induced very early; to avoid this, we used the pre-treatment stage as a control in both the phenotypic and the genetic analyses.

**Table 1:** Leaf and apex sampling times for flower mapping and gene expression analysis, expressed in terms of days after sowing and weeks of treatment.

| Date | Days after sowing | Week of treatment | Leaf sampling | Apex sampling | Phenotyping |
| --- | --- | --- | --- | --- | --- |
| 14.04. | 29dys | Control | x | x | x |
| 30.04. | 43dys | W0 | x |  | x |
| 07.05. | 50dys | W1 | x |  |  |
| 14.05. | 57dys | W2 | x |  |  |
| 20.05. | 63dys | W3 | x | x | x |
| 28.05. | 71dys | W4 | x |  |  |
| 04.06. | 78dys | W5 | x | x | x |

### 2.2. Assessment of flower bud initiation and flowering status

The total numbers of leaves, runners, and inflorescences on each plant selected for harvesting were recorded at each harvest. One additional plant per treatment and replicate was cut at the base, after which the leaves from the crown and the buds in each leaf axil were dissected under a stereo microscope (Figure 1B). Each meristem was classified as a branch crown (BC) or a runner and the developmental stage of the buds in the BC was rated using a scale adapted from Taylor et al. (1997) that ranges from 1 to 7, with a value of 1 indicating a vegetative state and values of 2-7 denoting generative states with increasing levels of floral differentiation. The position and stage of development of each bud was drawn schematically to create a ‘flower map’ in which the lower-most buds correspond to the oldest leaves (outermost in the plant), and the higher positions corresponded to the newest leaves (innermost buds in the crown). A typical ‘flower map’ is shown in Figure 1. This combination of methods made it possible to link genetic and phenotypic changes.

**Figure 1:**
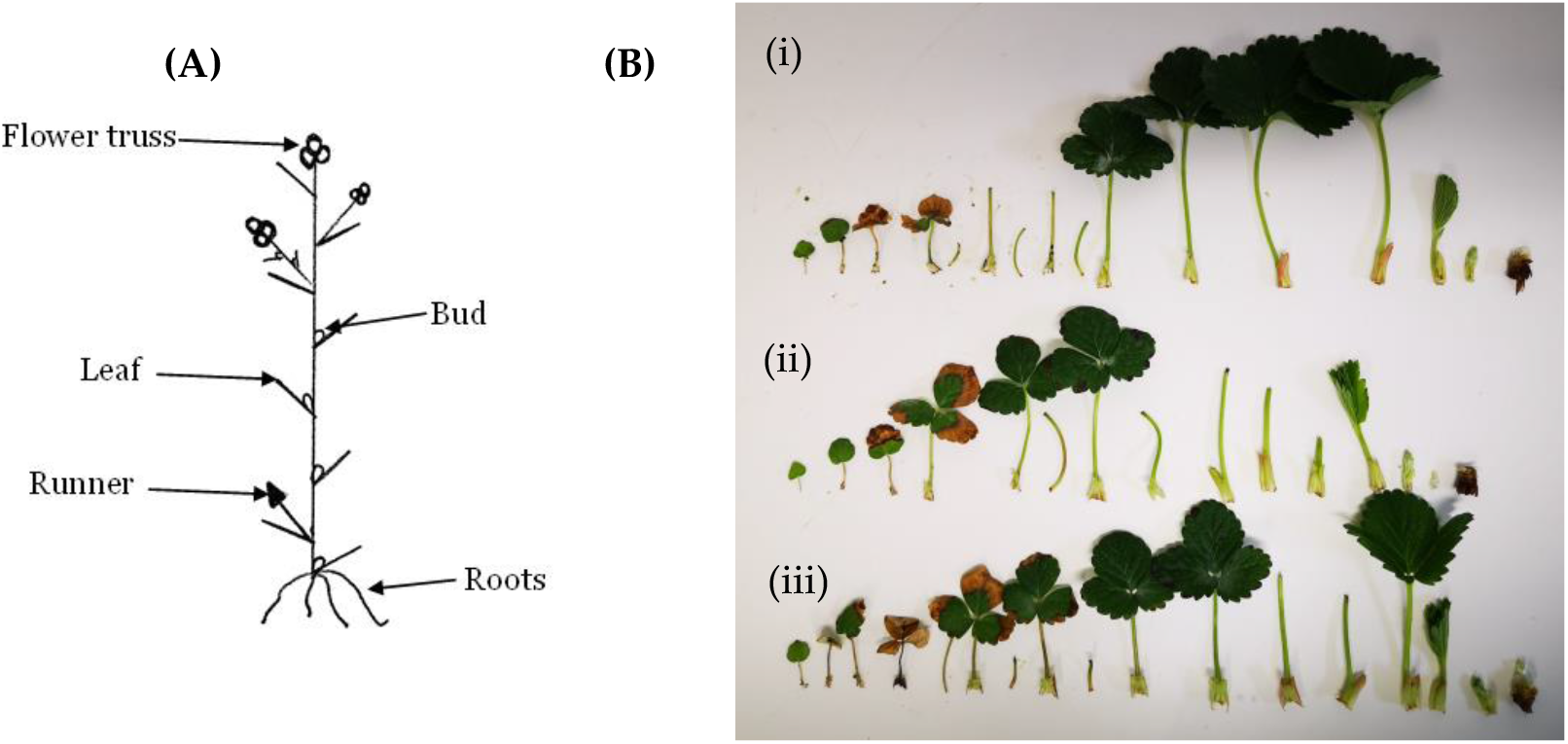
Schematic representation of the stage of development and position of buds within a branch crown (BC) of strawberry, also known as a ‘flower map’ (A), and the full set of dissected meristems from 3 replicate plants (i, ii, iii) grown for 5 weeks (W5) at 12°C under long day conditions (B).

An extra set of plants (3 replicates with 3 plants each) was grown for 39 days under each of the treatment conditions described above and then forced for 63 days in a green-house under LD conditions at a minimum temperature of 20°C for assessment of flowering status and development. Open flowers and runners were counted and removed weekly. At the end of the forcing period, the total number of leaves and crowns as well as the petiole length of the last developed leaf of each plant were recorded. This extra set of plants was grown to further differentiate the SD and LD responses in the EB cultivar.

### 2.3. RNA extraction and quantitative RT-PCR

Leaf samples for analysis of *FaFT* gene expression were first collected on April 14, and then on days 0, 7, 14, 20, 28, and 35 of treatment. Three biological replicate leaf samples were collected for each treatment; each sample consisted of 9 leaf discs (5 mm in diameter), 1 from each of the 3 leaflets of the last fully developed leaf of 3 plants. Shoot apex samples were collected for RNA extraction once during preconditioning (on April 14) and used as controls for all gene expression calculations. Additional samples were collected 3 and 5 weeks after the start of the treatments (Table 1). Leaf and apex samples for RNA extraction were collected at ZT = 4 ± 0.5 h at the time points specified in Table 1, then immediately frozen in liquid nitrogen and stored at -80°C.

RNA was extracted using a modified version of the pine tree method [43]. The purified RNA was then treated with Turbo rDNase (AM2238, ThermoFisher), according to the manufacturer’s recommendations. Reverse transcription of 500ng of total RNA was performed using the iScript cDNA synthesis kit (Bio-Rad Laboratories, Inc, California, USA) following the manufacturer’s instructions. Quantitative PCR was performed in 20 μl reaction mix containing 4 μl of 10-fold diluted cDNA, 2x DyNAmo Flash SYBR Green master mix (Thermo Scientific, Waltham, MA), and 0.5 μM primers. The Ct values of the test genes were normalized to the housekeeping gene *FaActin (FaACT)*, using results for the pretreated samples as controls. Quantifications were performed for nine biological replicates, with three technical replicates for each biological replicate. Relative gene expression was analyzed using the 2-ΔΔCT method. The RT-qPCR primers used in this study are listed in Supplementary Table A1.

### 2.4. Statistical analysis

ANOVA and regression analysis were conducted followed by Tukey’s multiple comparison tests for sub-factors (as stated under the figures). Spearman correlations between the expression of the genes analyzed in the apex (representing genotype data) and the meristem stage (MS, representing phenotypic data) were computed based on 3 biological replicates. All statistical analysis was done in R 4.0.3 [44].

## 3. Results

### 3.1. Effects of the environment on plant growth and development

The meristems were undifferentiated (Figure 2A) before the start of the treatment (Control and W0). At W3, the number of undifferentiated meristems was higher in SD than in LD except at 12°C (Figure 2A). Over the course of the experiment, significantly more meristems remained vegetative in SD than in LD. Although this decline occurred in all treatments, it was observed significantly earlier (W3) at 26°C than at lower temperatures.

**Figure 2:**
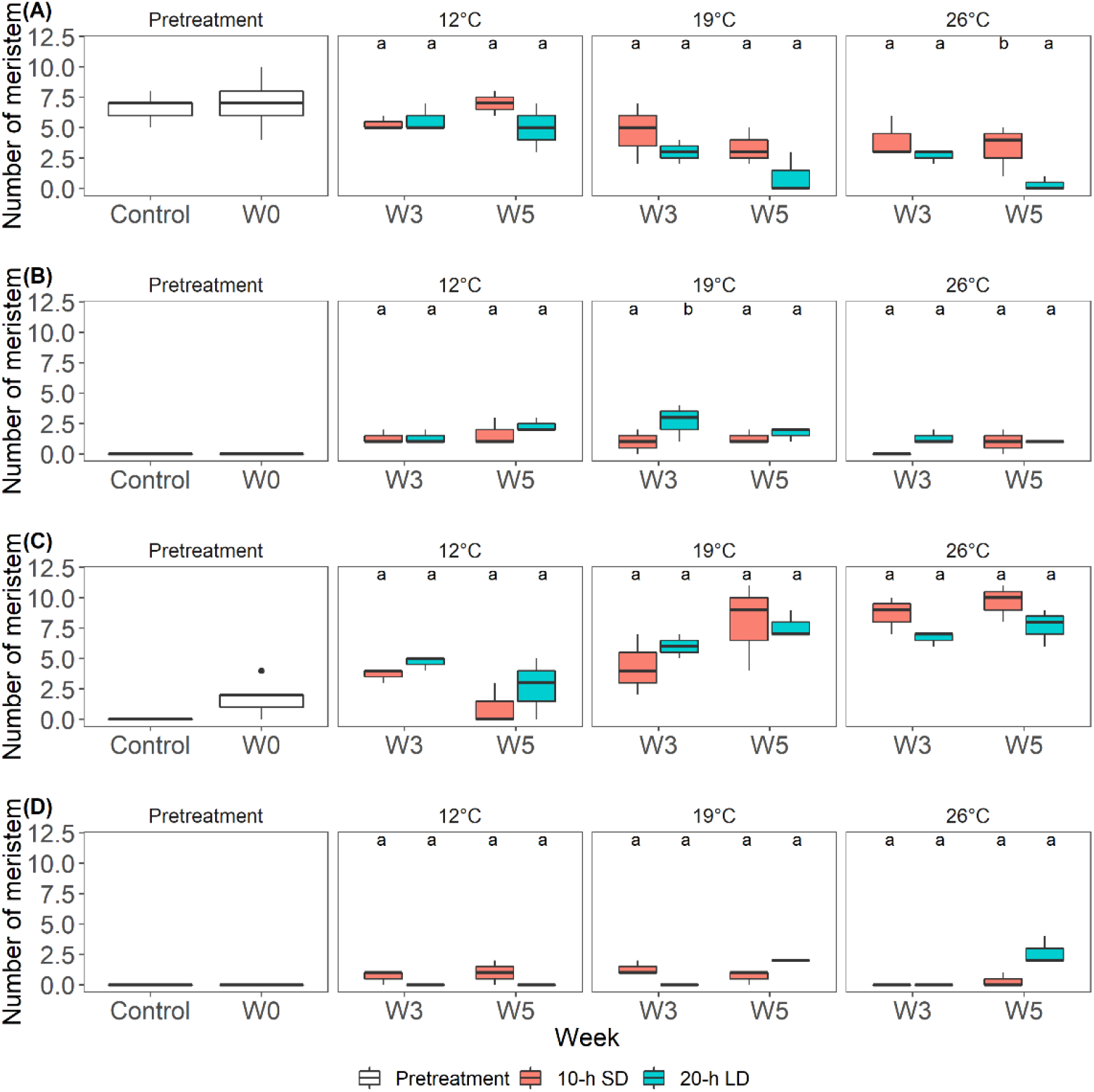
Average numbers of meristems scored as undifferentiated (A) and generative (B), and numbers of meristems that differentiated into runners (C) and branch crowns (BC) (D) during the 5-week treatment period (N=3 plants per treatment). Letters indicate significant differences between SD and LD photoperiods according to ANOVA and Tukey’s test (*p* <0.05). The ‘Control’ and ‘W0’ samples were collected before the start of the experiment (Pretreatment).

By W3, the number of generative meristems had increased in all treatments except those at 26°C SD, where the first generative meristem was observed in W5 (Figure 2B). The increase in generative meristems was initially photoperiod-independent at 12°C but the number of buds differentiating into generative meristems was significantly higher in LD than in SD at 19°C.

Runners (Figure 2C) started to appear at the start of the experiment (W0) and increased over the course of the 5-week treatment. The number of runners increased with the temperature, and there were slightly more runners in SD than in LD, although the photoperiodic effect was not statistically significant. From W3 onwards, the number of runners under SD conditions was significantly higher at 26°C than at lower temperatures. By W5, the number of runners at 12°C was significantly lower than at higher temperatures under both SD and LD conditions (Table A2).

Few BC were formed and their formation was delayed to W5 at 26°C. At 12°C, BC were observed in SD only. The same pattern was observed by W3 at 19°C, but the opposite trend was observed at this temperature by W5, when there were more BC in LD than in SD. The latter was also true at 26°C.

The meristems started differentiating at W0, when 16.1% of the meristems had differentiated into runners. By W3, the transition to generative development had started in all treatments except the 26°C SD treatment. By W5, some of the meristems in all of the treatments had become generative, and a higher percentage of generative meristems were observed in LD than in SD except at 26°C. At 19°C and 26°C, around 50% of the meristems differentiated into runners, independently of the photoperiod. From W3 to W5, an increasing percentage of meristems became runners rather than generative meristems in LD at 19°C and 26°C. The proportion of BC was highest in W5 under LD conditions at 19°C and 26°C, while no BC was observed in LD at 12°C (Figure 3).

**Figure 3:**
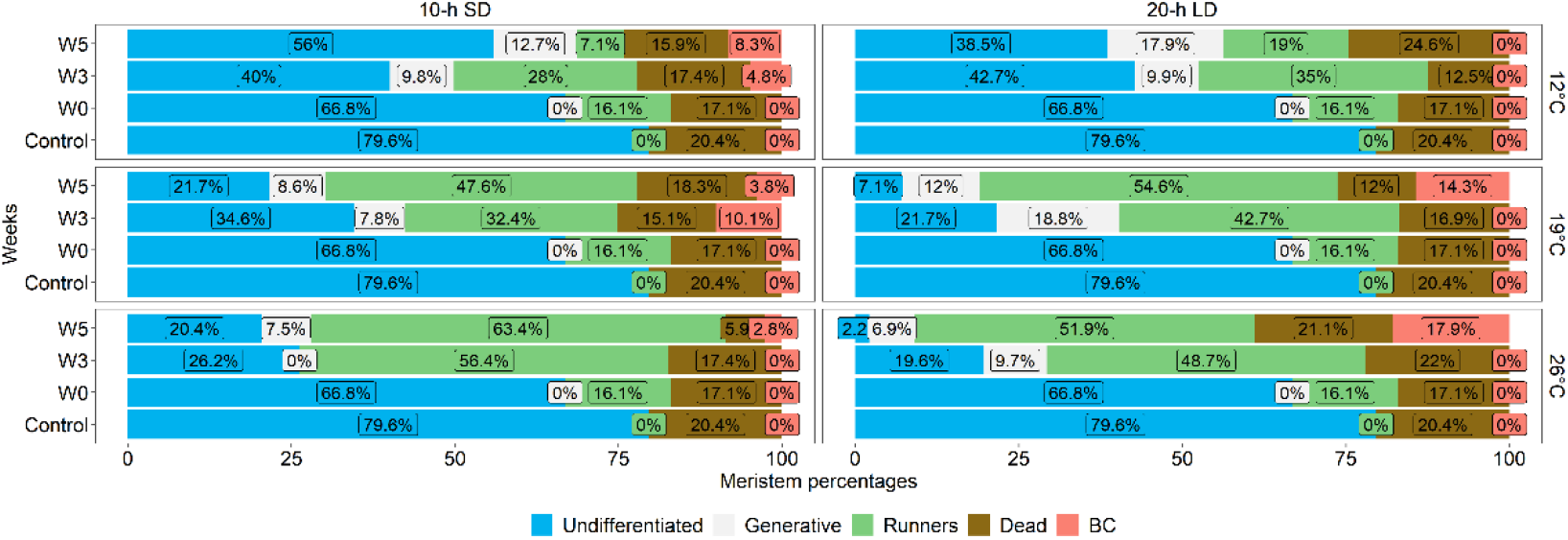
Average percentage of meristems that were vegetative (blue), generative (white), differentiated into runners (green), dead (brown), and differentiated into branch crowns (BC, pink) at W0 (n=9 plants), W3 (n=3 plants), and W5 (n=3 plants) during the experiment. Control data (representing n=9 plants) were collected before the start of the experiment (Table 1).

Meristems assigned scores ≥2 were considered generative. The results presented in Figure 4 show that the plants in LDs were always at a more advanced developmental stage than their SD counterparts in all treatments. By W3, the plants were generative in all treatments except in SD at 26°C. Moreover, there was considerable variation in meristematic development under SD conditions even after 5 weeks (W5), with plants ranging from vegetative to generative. Conversely, all plants in LD treatments were generative by week 5. The transition to generative development at 26°C was significantly delayed in SD when compared to LD at both W3 and W5.

**Figure 4:**
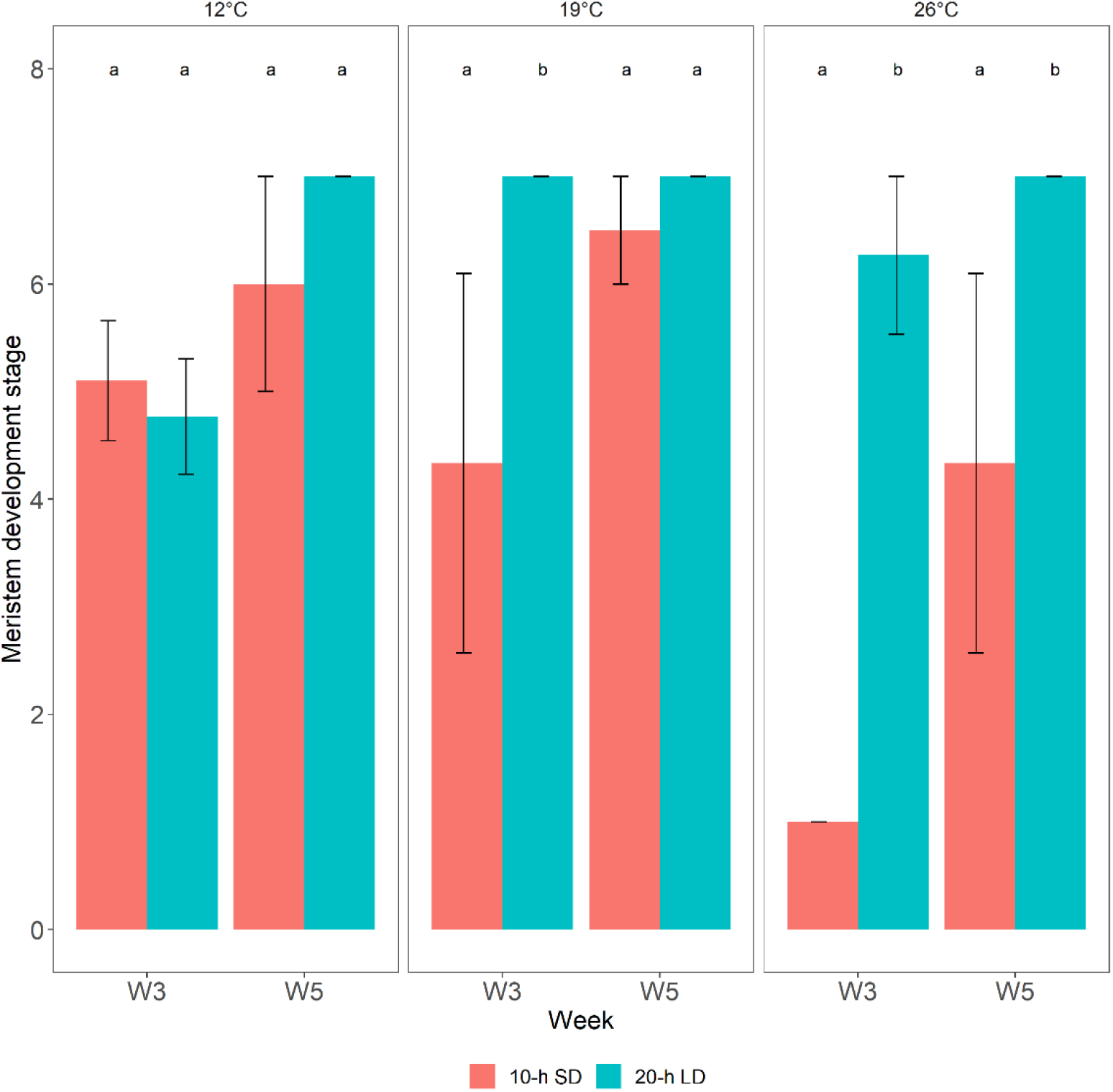
Average floral development stages (n=3) at temperatures of 12°C, 19°C, and 26°C under 10-h SD and 20-h LD conditions. Before initiating the treatments, all meristems were vegetative (stage=S1; data not shown). Floral stages were evaluated using Taylor’s 7-stage scale (1997) in which stage 1 (S1) denotes vegetative meristems, stage 2 (S2) corresponds to the first visible evidence of flower initiation, and stage 7 (S7) denotes fully differentiated meristems in which primordia of all floral organs are visible. Error bars represent standard errors of means; letters indicate significant differences between SD and LD photoperiods according to regression analysis and Tukey’s test (*p* <0.05).

### 3.2. Post treatment plant development

The appearances of representative plants after 39 days under the tested growing conditions are shown in Figure 5. At this stage, all plants were subjected to forcing for 63 days in a greenhouse maintained at 20°C under 20-h LD conditions. This was the point at which the first inflorescences became visible in plants grown in LD at 19 and 26°C.

**Figure 5:**
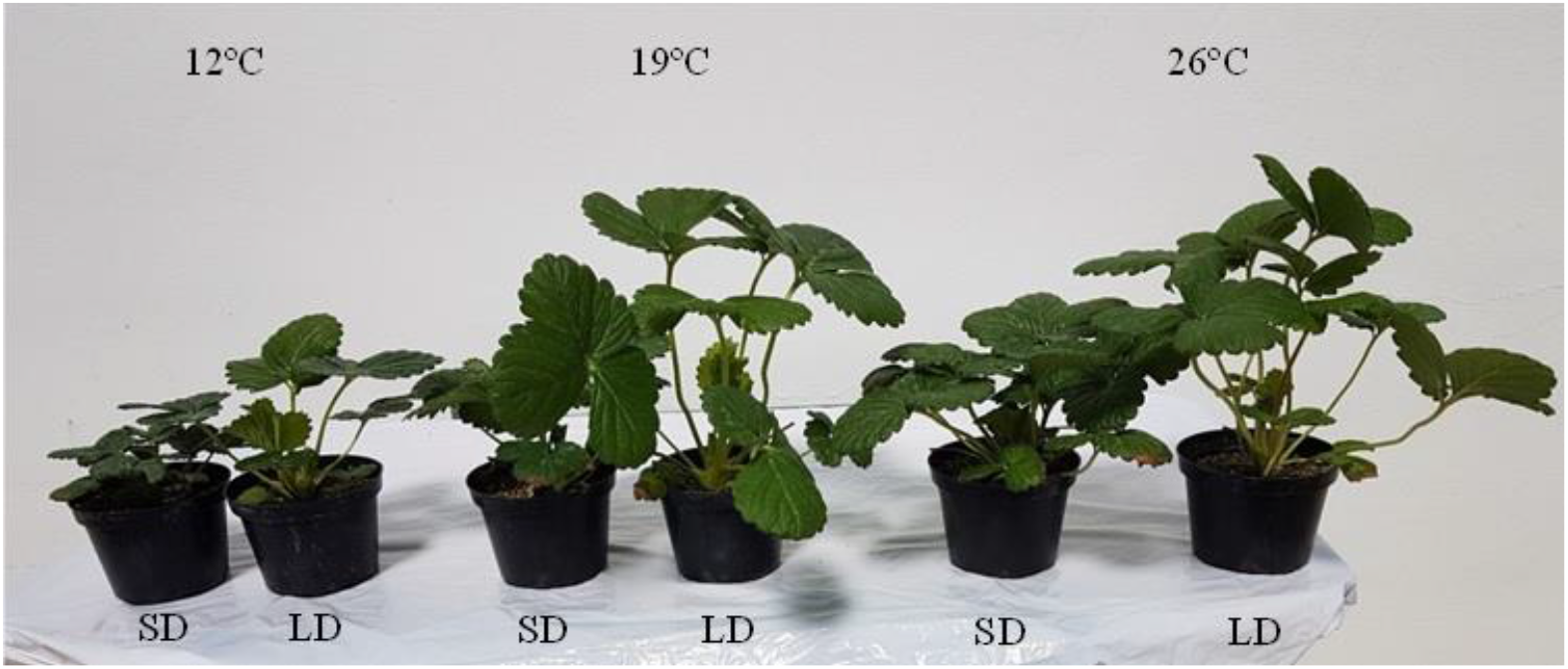
Appearance of representative strawberry cv. Delizzimo plants after 39 days of treatment under the indicated conditions; SD denotes 10-h SD and LD denotes 20-h LD.

Time courses showing the weekly cumulative appearance of runners and open flowers during the 63-day forcing period are shown in Figure 6. Runner production increased with the preconditioning temperature and was enhanced by SD conditions at all temperatures. Flowering also increased with the preconditioning temperature, but was enhanced by LD at all temperatures, unlike runnering (Figure 6). It should also be noted that flowering was strongly delayed in plants preconditioned under SD conditions at 26°C, indicating that not all of these plants had been initiated before forcing.

**Figure 6:**
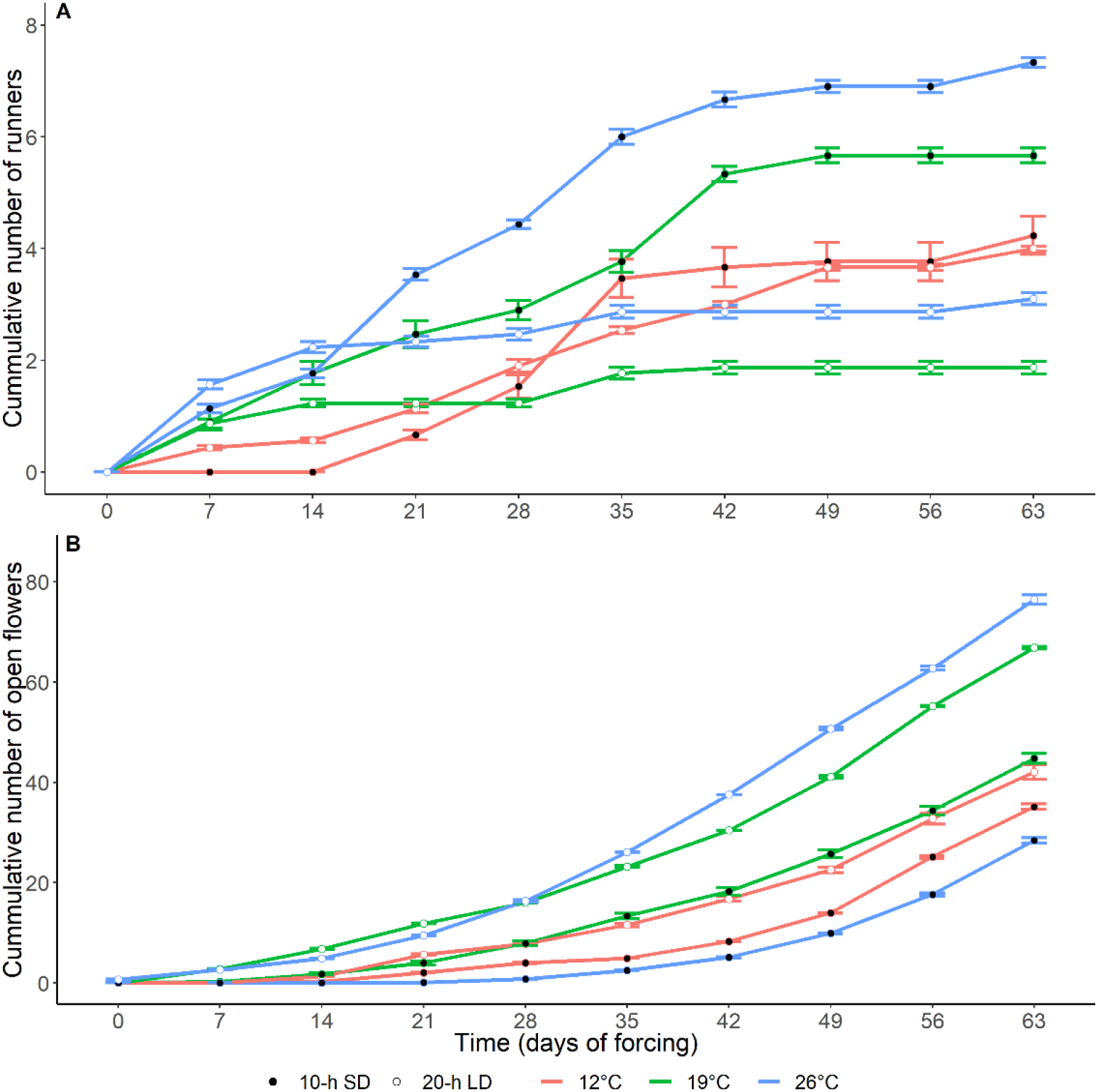
Cumulative increases in the numbers of runners (A) and open flowers (B) over the 63-day forcing period in ‘Delizzimo’ strawberry plants (n = 3) preconditioned under different temperature and photoperiodic conditions. Values are means ± standard errors of three replicates of five plants each.

### 3.3. FaFT1 expression in leaves

*FaFT1* expression was undetectable in leaves before the start of treatment (Control) and at W0 (Table 1) but was upregulated in LD by W1 at all temperatures (Figure 7). Its expression generally peaked by W2 and then declined again. Under SD conditions, however, *FaFT1* expression was very weak or undetectable on all sampling dates.

**Figure 7:**
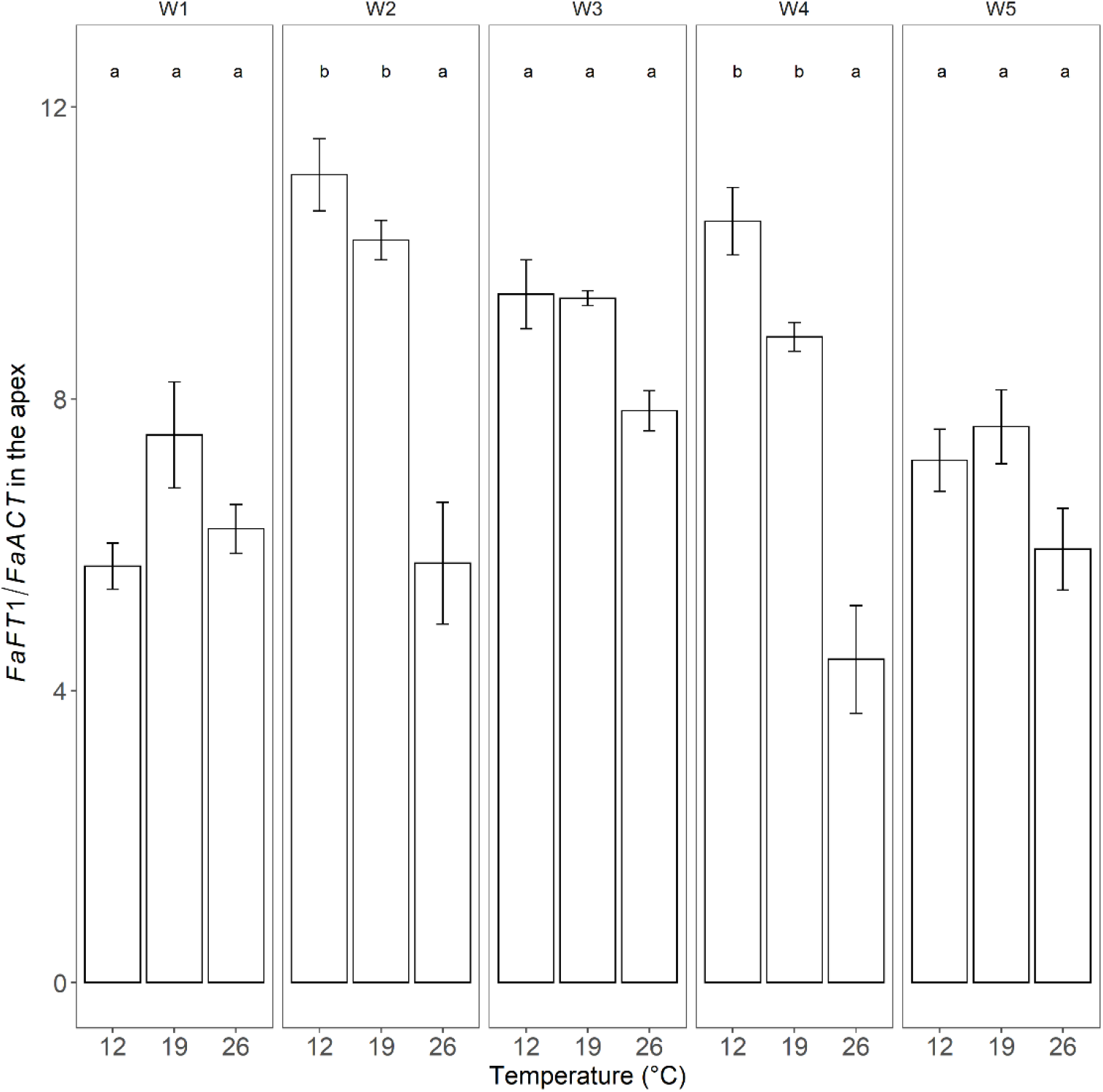
The weekly relative expression of *FaFT1* in the leaves of the strawberry cv. Delizzimo grown at 12°C, 19°C, and 26°C under LD conditions for 5 weeks. The plotted results represent average expression levels in three biological replicates and three technical replicates. Relative expression was normalized to the housekeeping gene *FaACT* as an internal standard and pre-treatment expression was used as a control (Table 1). Error bars represent standard errors of means and letters indicate significant differences between the temperature treatments based on logistic regression and Tukey’s test (*p* <0.05). Three biological replicates were collected, each comprising 9 leaf discs (5 mm in diameter), one from each of the three leaflets of the last fully opened leaf.

### 3.4. Effect of photoperiod and temperature on flowering genes in the apex

We also studied the expression of known flowering genes in apex samples taken at the time points indicated in Table 1. *FaSOC1* is downstream of *FaFT1*; its expression was significantly higher in LD than in SD at all temperatures (Figure 8A) and correlated with that of *FaFT1* (Figure 7). Additionally, *FaSOC1* expression was significantly lower in SD than LD and was also lower at 26°C than at 12°C (Figure 8A). Its expression also decreased significantly from W3 to W5 in plants grown in LD at 12°C and 26°C (Table A3).

**Figure 8:**
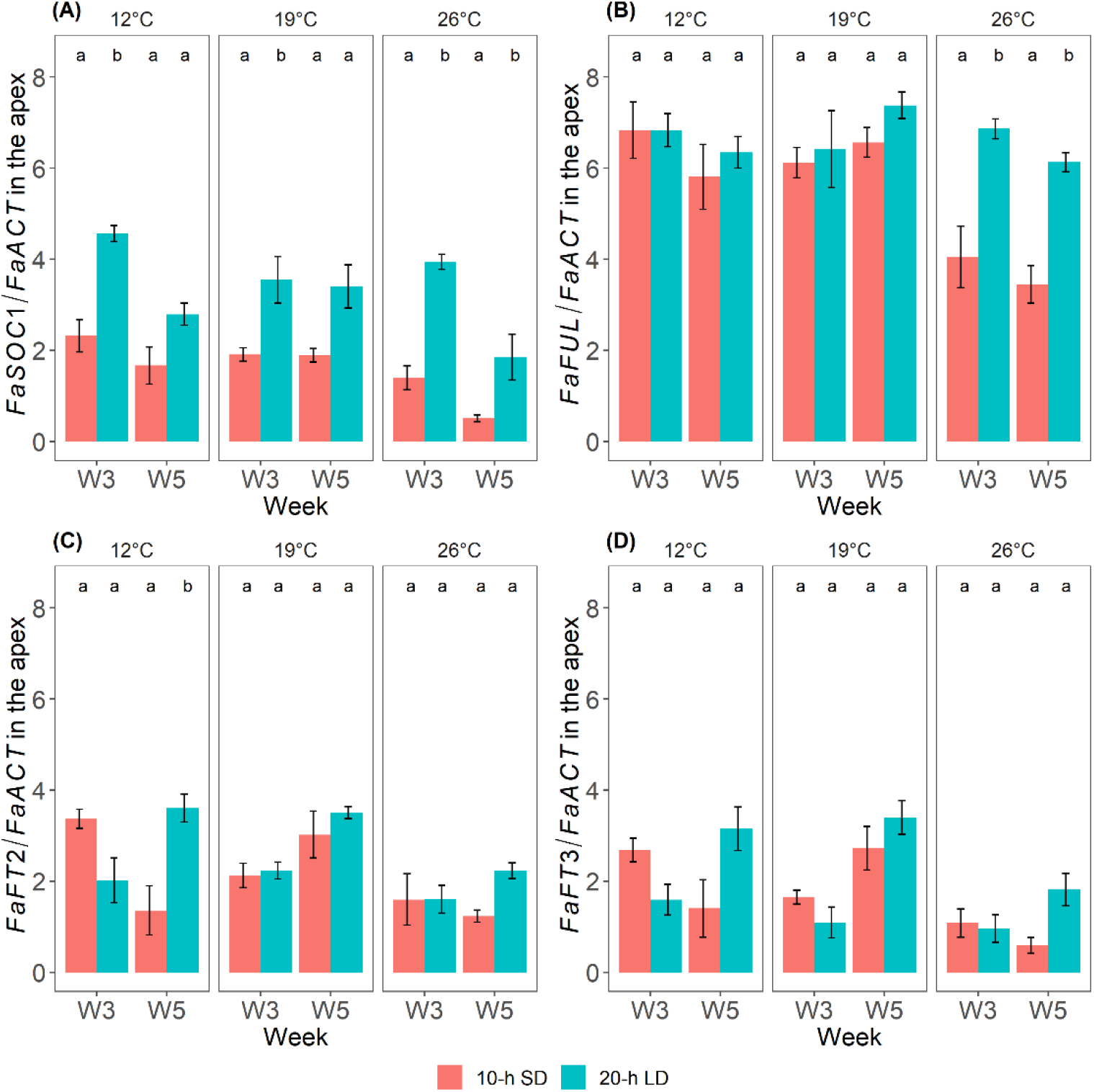
Relative expression of *FaSOC1* (A), *FaFUL* (B), *FaFT2* (C) and *FaFT3* (D) in the apices of strawberry cv. Delizzimo plants grown at 12°C, 19°C, and 26°C under SD and LD conditions for a period of 5 weeks. The data are average expression levels of three biological and technical replicates. The relative expression was normalized to the housekeeping gene, *FaACT* as an internal standard, using pre-treatment data as a control (Table 1). Error bars represent standard errors of means and letters indicate significant differences between LD and SD photoperiods evaluated using a linear mixed effects regression model and Tukey’s test (*p* <0.05).

To evaluate flower bud initiation, we examined the expression of *FaFUL*, a meristem identity gene previously identified as a floral promoter [24], as well as *FaFT2* and *FaFT3*, which were previously used as markers of flower initiation in SF Japanese cultivars [17,25,27]. *FaFUL* was highly expressed in all treatments (Figure 8B) except SD at 26°C. By W3, both *FaFT2* (Figure 8C) and *FaFT3* (Figure 8D) were upregulated. *FaFT2* was initially expressed more strongly under SD than LD conditions at 12°C and the same was true for *FaFT3*, although the difference was not statistically significant in the latter case (Table A4). Both *FaFT2* and *FaFT3* were weakly expressed at 26°C, but the expression of both genes increased from W3 to W5 under LD conditions.

## 4. Discussion

### 4.1. Environmental effects on general plant growth and development

The strawberry meristem can remain undifferentiated or develop into runners, BC, or generative meristem. Meristem phenotypes were categorized as vegetative or generative based on an adapted variant of the 7-stage scale proposed by Taylor et al. (1997), where stage 1 denotes a vegetative meristem and stages 2-7 denote progressively more advanced stages of generative development [19]. Although runners are also vegetative, we distinguished them from vegetative meristems to avoid disproportionately inflating the number of vegetative meristems. Our objective in this work was to simultaneously analyse meristem phenotypes and the expression of known flowering genes to compare traditional method of evaluating meristem development to approaches based on molecular analysis of meristematic changes.

The studied cultivar exhibited the typical features of a quantitative LD plant with earlier flowering in LD than in SD at all temperatures, and especially at 26 °C (Figure 4). Runner formation, on the other hand, was significantly enhanced by high temperature (Figure 2C, Figure 3, and Table A2). However, young strawberry seedlings have a short juvenile period during which runner formation occurs independently of the environment, so-called “juvenile runnering” [20]. Therefore, runner formation in ‘Delizzimo’ was initially unaffected by the photoperiod. However, after termination of the treatments (W5), there was a shift to SD-stimulation of runner formation that was associated with flower initiation in LD. This negative relationship between flowering and runnering in adult plants is clearly illustrated in Figure 6, which shows the after-effects of the 5-week treatments. Since a meristem can differentiate into only one organ, the results reveal a clear inverse relationship between flowering and runnering. Runner formation was always highest in plants grown under SD conditions, while flowering was highest in LD, and increased with temperature in both cases. Figure 6 also shows that while plants grown in LD at 19 and 26°C reached anthesis after 7 days of forcing, this stage was reached only after 35 days in plants initially grown under SD conditions at 26°C. Some plants grown under the latter conditions had not even initiated flowers before being transferred to LD forcing (Figure 4).

### 4.2. Effects of the environment on FaFT1 expression

To study induction in EB strawberry plants, we analyzed *FaFT1* expression in the leaves of ‘Delizzimo’ seedlings grown in SD and LD at 12°C, 19°C, and 26°C, using the primers designed by Koskela et al. (2016) (Table A1). Our results were consistent with those previously reported by Nakano et al. (2015) and Koskela et al. (2016) for SF cultivars: *FaFT1* expression was high under LD conditions but not detectable under SD conditions. This confirmed the LD flowering trait in EB strawberry [2] and demonstrates that floral induction in ‘Delizzimo’ leaves is mediated by *FaFT1* upregulation. This upregulation started as early as W1 in all treatments, showing that the plants were induced very early in the experiment (Figure 7). Early induction in the EB strawberry has also been previously reported, leading to some misinterpretation of results. To avoid this problem, gene expression was also measured before initiating treatment to obtain control data [10,12,42].

*FaFT1* expression was initially highest at 19°C and was high at both 12°C and 19°C in most treatment weeks, with lower expression at 26°C. However, its expression declined steadily at all temperatures from W2 to W5.

### 4.3. Post-induction floral bud initiation

Although floral induction cannot be deduced by flower mapping, our results show that if flower mapping is performed at an appropriate time, its results are consistent with those obtained by analysing molecular markers related to flowering (Figures 4, 7, and 8). Although the method is labour-intensive and requires special training, it remains essential for breeders that rely on its results to make important decisions concerning flowering, and thus yield [15,45,46]. We also see that even when flowering has been induced, differences in post-induction environmental conditions can still influence initiation and subsequent stages of floral development (Figure 6), in accordance with previous reports [47,48]. To circumvent these problems, it is necessary to either perform multiple rounds of flower mapping or to use molecular markers.

The present results also indicate that the *FaSOC1, FaFUL, FaFT2*, and *FaFT3* genes of the photoperiodic flowering pathway interact in mediating floral initiation in the apex. By W3, all plants were generative irrespective of treatment, except those in SD at 26°C, which were still mostly vegetative (Figure 4). All plants grown under LD conditions had been induced by W1 (Figure 2), which coincided with clear upregulation of *FaFT1* (Figure 7). Although *FaFT1* expression not detectable in the leaves of plants grown under SD conditions, upregulation of *FaSOC1* (which is downstream of *FaFT1*) confirmed floral initiation by W3 in plants grown under both LD and SD conditions and at all temperatures (Figures 4 and 8). In other words, induction in the leaves was LD-dependent in ‘Delizzimo’, whereas floral initiation in the apices was photoperiod-independent.

Koskela et al. (2016) showed that silencing of the floral repressor (antiflorigen) *FaTFL1* caused photoperiod-independent perpetual flowering in the SF cultivar ‘Elsanta’. They also provided evidence that *FaTFL1* is generally expressed only under LD conditions in cultivated strawberry plants, whereas *FaSOC1* is expressed in both SD and LD. Similar results were obtained in an EB *F. vesca* genotype by Mouhu et al. (2013). Furthermore, Sønsteby and Heide (2008) found that the flowering responses of seed-propagated EB *F. vesca* cultivars were identical to those of the octoploid EB F1-hybrid ‘Elan’, a close relative of ‘Delizzimo’.

Lei et al. (2013) isolated and characterized *FaSOC1* in the SF cultivar ‘Camarosa’, by studying gene expression in the apex under SD floral induction conditions at 18°C/15°C for 6 weeks. They found that *FaSOC1* expression increased in the apex during the first 2 weeks under SD inductive conditions in the SF cultivar but then fell dramatically [49]. We observed a similar decrease in *FaSOC1* expression between W3 and W5 under LD conditions (Table A3), which was statistically significant at 12°C and 26°C but not at 19°C. The physiological implications of this decline are currently unknown.

Nakano et al. (2015) detected no significant differences in gene expression between photoperiodic treatments in the Japanese SF cultivar ‘Nyoho’. However, studies on three SF cultivars by Koskela et al. (2016) showed that *FaSOC1* expression is cultivar-dependent at 18°C. Specifically, under LD conditions its expression decreased after two weeks but then eventually increased in ‘Honeoye’, whereas in ‘Alaska Pioneer’ and ‘Polka’ it decreased steadily over time. However, *FaSOC1* expression in all three cultivars increased under LD conditions following 6 weeks of SD growth, showing that *FaSOC1* was upregulated by delayed LD exposure. This suggests that SF strawberry cultivars are in fact SD-LD plants [3,7]. Koskela et al. (2016) also performed a temperature experiment with the SF cultivars ‘Glima’ and ‘Elsanta’, revealing that *FaSOC1* expression is temperature-sensitive. Prolonged exposure to low (9°C) and high (21°C) temperatures resulted in a lower expression than at 15°C (Table A3). However, Rantanen et al. (2015) found that the photoperiod and temperature flowering pathways converge at *FvSOC1* in SF *F. vesca*. Keeping in mind the strong interaction between photoperiod and temperature in the control of flowering in both diploid and octoploid strawberries [3], this convergence would be expected to have complex effects on *FaSOC1*, which may partially explain its divergent expression patterns in different genotypes. Liang, et al (2022) also found a decline in *FaSOC1* levels during the start of the floral initiation followed by an increase in the expression 2 weeks later [41]. However, the increase during their experiment could also be due to the fluctuations in the temperature and photoperiod in the greenhouse over the course of the experiment. This further strengthens our claim that changes in the environment can affect the genes as well as the architecture of the strawberry even after induction.

Taking all evidence into account, it seems that *FaSOC1* expression may be cultivar-dependent. However, its expression was higher in LD than in SD in all cultivars at all temperatures, and was increased by moving the plants from SD to LD conditions [24]. It was previously suggested that *FaSOC1* may be involved in runner formation, which was subsequently confirmed in the woodland strawberry [37]. However, since post-juvenile runner formation in ‘Delizzimo’ was inversely related to flowering in this study, runner formation was promoted by SD because of the indirect effect of LD-induced flowering.

We also studied the expression of the floral meristem identity gene *FaFUL*, revealing that it was upregulated in all treatments by W3 (Figure 8). The timing of its upregulation thus coincides with floral initiation, making *FaFUL* a potentially valuable early indicator of floral initiation in the EB ‘Delizzimo’ strawberry.

### 4.4. Environmental effects on FaFT2 and FaFT3 expression

Nakano et al. (2015) proposed that *FaFT2* and *FaFT3* function as florigens in the Japanese SF cultivar ‘Nyoho’ in SD at cooler temperatures while *FaFT1* has this function in LD at warmer temperatures. However, recent reports on the basis of transcriptomics data have presented conflicting results, where Koembuoy et al. (2020) identified *FaFT3* as a floral inducer for Japanese SF cultivars; while, Liang et al. (2022) detected an increase in the *FaFT2* expressions in the apex, during floral initiation in ‘Benihoppe’, a SF cultivar [41]. Perrotte et al. (2016) on the other hand, suggested that the strawberry homolog *FaFT2* may be responsible for the everbearing character in octoploid strawberry [50].

We therefore studied the expression of *FaFT2* and *FaFT3* using the primers developed by Nakano et al. (2015) and found that both genes were upregulated independently of temperature and photoperiod. We also found that their expression was initially higher in SD than in LD at 12°C, and that they exhibited comparatively low expression at 26°C (Figure 8). However, their expression in the apices in both SD and LD suggests that they play a role in the downstream floral initiation process by interacting with other gene members of the photoperiod pathway.

We could not determine whether all of the studied genes act together to trigger initiation, or whether each individual gene has a specific threshold level of expression that is required for initiation. However, there were significant positive correlations between the expression levels of all the genes (Figures A1 and A2). Their expression was also positively correlated with meristem stage advancement, and this correlation became stronger at higher temperatures in the apex. In addition, there was a positive correlation at lower temperatures for *FaFT2* and *FaFT3* (Figures 8, A1 and A2). Further functional studies are needed to robustly characterize the roles of these genes in EB cultivars, however, because the limited sample sizes used in this work, the variation between replicates, and other limitations of the experiment made it difficult to draw reliable conclusions about the specific functions of the various flowering-related genes.

While flower induction and initiation are poorly understood in the perpetual-flowering garden strawberry, they are better understood in the perpetual-flowering H4 mutant wood strawberry that does not express a functional *FvTFL1* homolog [24,31,32]. In this mutant, the floral repressor, *FvFT1* is strongly expressed in the leaves under LD flower-inducing conditions, in exactly the same way as we found in ‘Delizzimo’. A possible explanation is that the EB gene also functions as a floral repressor. This possibility is supported by the identical LD flowering responses seen in perpetual-flowering *F. vesca* genotypes and the ‘Elan’ garden strawberry F1 hybrid over a wide range of temperatures [51] and should therefore be investigated further.

## 5. Conclusions

In summary, our experimental results showed that the flower mapping method used by breeders is reliable if the environment is consistent because development of floral structures can be delayed by changing environmental cues even after initiation. Characterization of the genotypic and phenotypic features of the seed-propagated everbearing F1 hybrid ‘Delizzimo’ revealed that the floral meristem identity gene *FaFUL* is an early indicator of floral initiation in EB strawberry. However, further studies are needed to investigate the functional relationships among the FT homologs and genes acting downstream of them. Finally, we recommend the use of F1 hybrid seedlings in studies of this kind because they provide plant material that is genetically identical and, crucially, not biased by previous flowering history.

## Supplementary Materials

The following supporting information can be downloaded at: www.mdpi.com/xxx/s1, Figure S1: title; Table S1: title; Video S1: title.

## Author Contributions

S.S. was responsible for conceptualizing and project design, visualizing the data, writing the original draft, review and editing. R.R. was responsible for the methodology design of the physiological studies, visualizing the data and contributed to the writing, review, and editing. PBK. was responsible for the methodology design of the gene analysis and analyzed the gene expression data and contributed to the methodology writing. R.V. was responsible for project administration and supervised P.B.K. O.M.H. supervised R.R. and contributed to the writing, review, and editing. A.S. was responsible for conceptualizing, methodology design, supervised R.R., and contributed to the writing, review, and editing. S.K. was responsible for resources, supervision of S.S., project administration, and funding acquisition.

## Acknowledgments

We acknowledge the help and support offered by Dr. Sharon Hill by providing lab space for parts of the molecular work conducted in this study. We would also like to acknowledge Salla Marttila for assisting with acquiring liquid nitrogen for the RNA extraction process.

## Appendix A

**Table A1:**
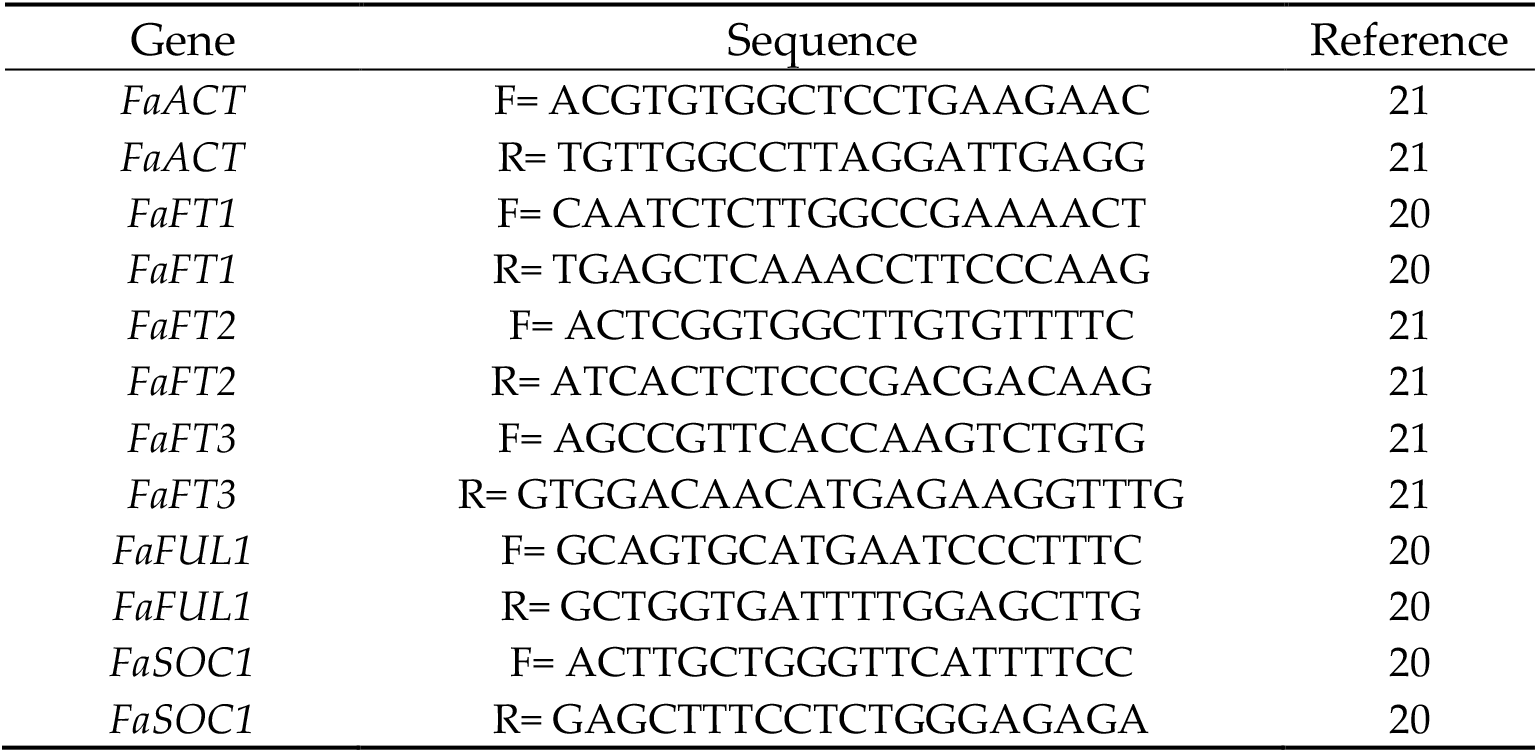
qPCR Primers used in the study

**Table A2:** Statistical significance of differences between the number of meristems that differentiated into runners in the different temperature treatments (12°C, 19°C, and 26°C). The grouping is based on Holm’s test; treatments with the same letters do not differ significantly (α = 0.05).

| Week | Temperature (°C) | Photoperiod | Phenotype | Holm's test |
| --- | --- | --- | --- | --- |
| W3 | 12 | SD | Runners | a |
| W3 | 19 | SD | Runners | a |
| W3 | 26 | SD | Runners | b |
| W5 | 12 | SD | Runners | a |
| W5 | 19 | SD | Runners | b |
| W5 | 26 | SD | Runners | b |
| W5 | 12 | LD | Runners | a |
| W5 | 26 | LD | Runners | b |
| W5 | 19 | LD | Runners | b |

**Table A3:**
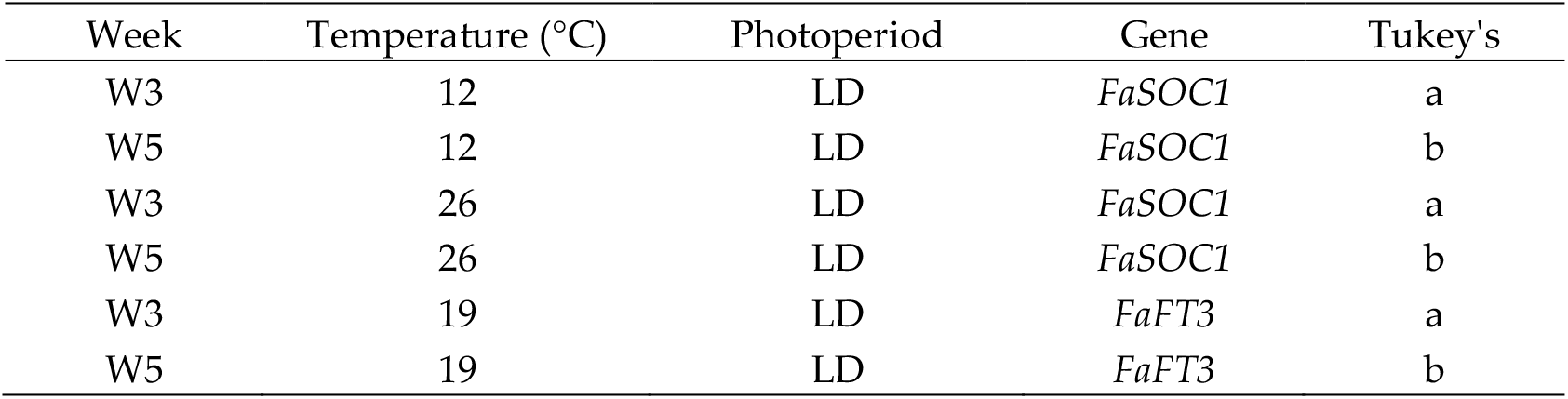
Summary of a mixed effects regression analysis of the expression of the studied genes in W3 and W5. Results are only shown for treatments with significant differences in expression (α = 0.05) between the two time points according to Tukey’s test. The level of expression does not differ significantly between treatments labelled with the same letter in the right-hand column.

| Week | Temperature (°C) | Photoperiod | Gene | Tukey's |
| --- | --- | --- | --- | --- |
| W3 | 12 | LD | *FaSOC1* | a |
| W5 | 12 | LD | *FaSOC1* | b |
| W3 | 26 | LD | *FaSOC1* | a |
| W5 | 26 | LD | *FaSOC1* | b |
| W3 | 19 | LD | *FaFT3* | a |
| W5 | 19 | LD | *FaFT3* | b |

**Table A4:** Summary of a mixed effects regression analysis of the expression of the studied genes in the different temperature treatments (12°C, 19°C, and 26°C) in each week of the study. Results are only shown for samples and treatments exhibiting significant differences (α = 0.05) according to Tukey’s test. The level of expression does not differ significantly between treatments labelled with the same letter in the right-hand column.

| Temperature (°C) | Week | Photoperiod | Gene | Tukey's |
| --- | --- | --- | --- | --- |
| 12 | W5 | SD | *FaFUL* | ab |
| 19 | W5 | SD | *FaFUL* | a |
| 26 | W5 | SD | *FaFUL* | b |
| 12 | W5 | SD | *FaFT3* | ab |
| 19 | W5 | SD | *FaFT3* | a |
| 26 | W5 | SD | *FaFT3* | b |

**Figure A1:**
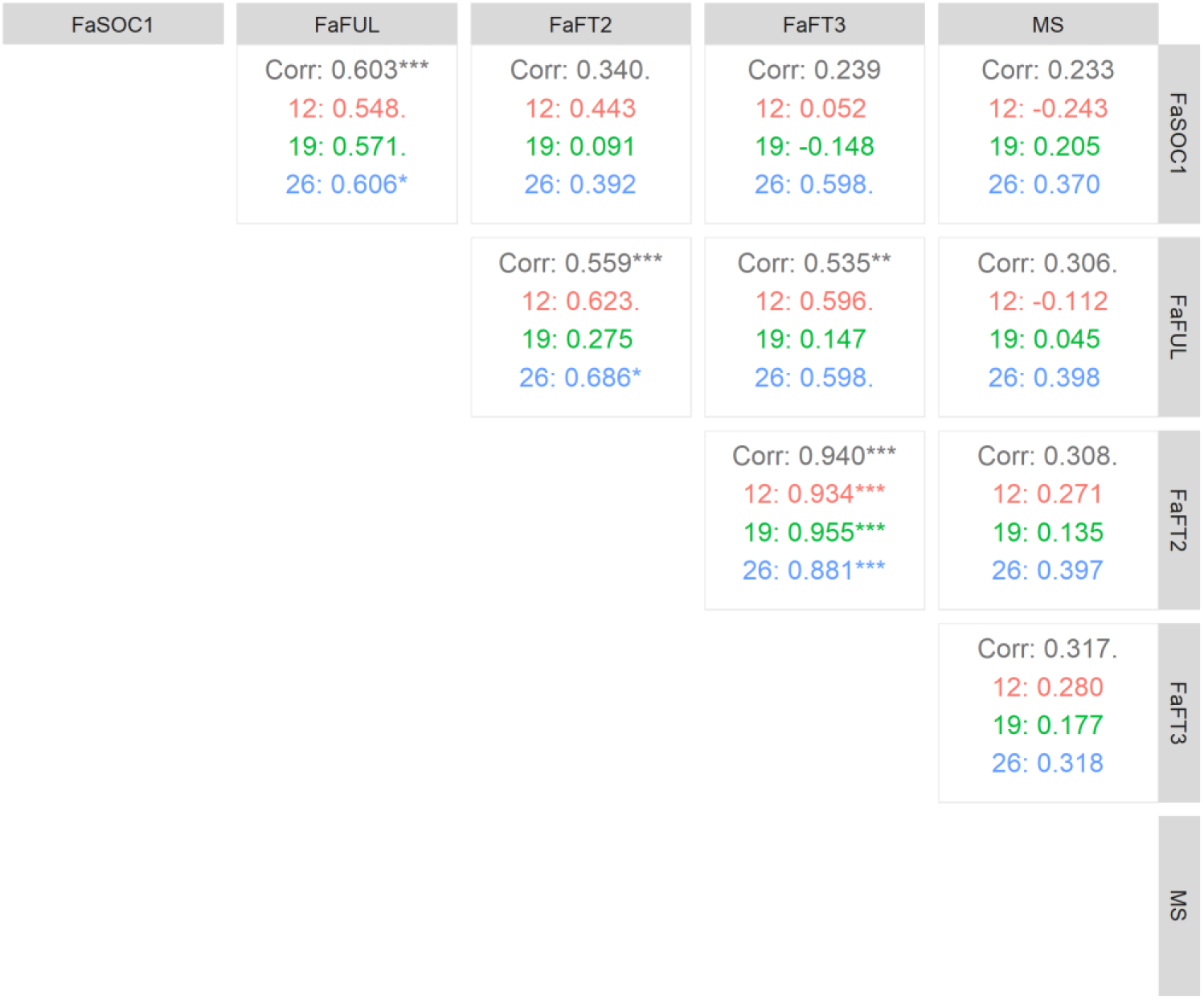
Spearman correlation coefficients between gene expression (*FaSOC1, FaFUL, FaFT2, FaFT3*) and meristem stage (MS; Figure 4) under the different temperature treatments (12ºC, 19ºC and 26ºC). Significance is indicated as follows: ‘***’ *p* < 0.001, ‘**’*p* <0.01, ‘*’*p* <0.05 and ‘.’ *p* < 0.1

**Figure A2:**
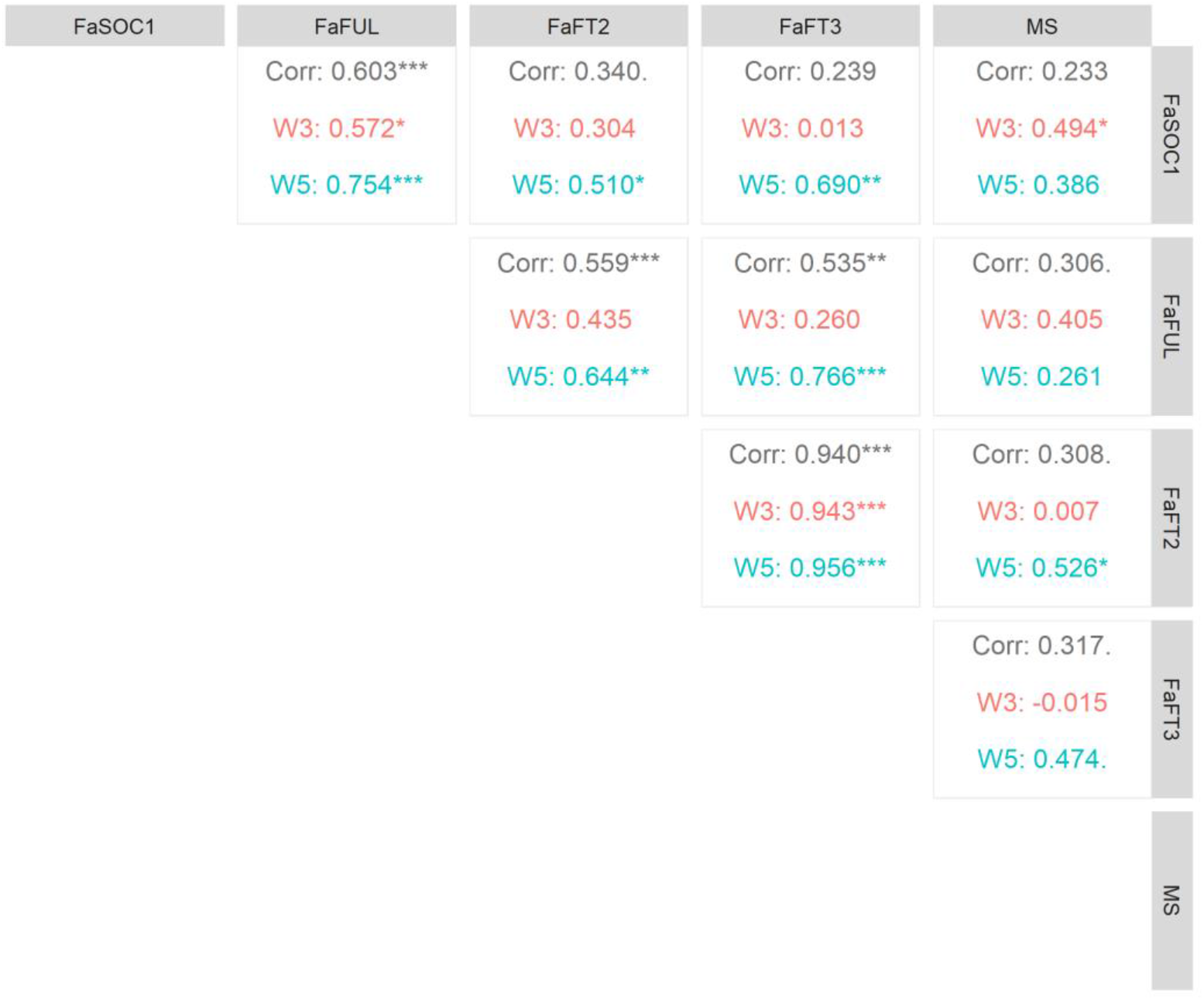
Spearman correlation coefficients between gene expression (*FaSOC1, FaFUL, FaFT2, FaFT3*) and meristem stage (MS; Figure 4) under SD and LD conditions. Significance is indicated as follows: ‘***’ *p* < 0.001, ‘**’*p* <0.01, ‘*’*p* <0.05 and ‘.’ *p* < 0.1

